# Bacterial-Mimetic Membranes Reveal Hopanoid-Associated Domain-Like Lateral Heterogeneity

**DOI:** 10.64898/2026.11.08.763706

**Authors:** Mateusz Rzycki, Maxime Kucharski, Natalia Szulc, Dominik Drabik

**Affiliations:** Department of Biomedical Engineering, Wroclaw University of Science and Technology, Wroclaw, Poland; Department of Physics and Biophysics, Wroclaw University of Environmental and Life Sciences, Wroclaw, Poland

**Keywords:** lipid membranes, microdomains, bacterial mimicking membrane, Giant unilamellar vesicle, molecular dynamics

## Abstract

Bacterial membranes are increasingly recognized as laterally heterogeneous assemblies whose local organization can regulate transport, stress adaptation, and interactions with membrane-active compounds. However, the physical principles by which bacterial lipid composition generates spatially distinct membrane environments re- main poorly understood, partly because experimentally tractable models that reproducibly exhibit lateral segregation and permit quantitative characterization are limited. Here, we show that a multicomponent, hopanoid-containing bacterial-inspired mem- brane spontaneously self-organizes into laterally distinct physical environments that are consistently resolved in experiments and molecular simulations. Giant unilamellar vesicles prepared from a lipid mixture inspired by lipid classes found in hopanoid- producing bacteria, including Methylorubrum extorquens, exhibit stable micron-scale lateral segregation, which is independently reproduced in compositionally matched coarse-grained molecular dynamics simulations. The resulting membrane environments differ in composition, packing, order, mobility, and mechanical properties. Hopanoid- associated domain-like regions are more condensed, more ordered, and less mobile, whereas PE-rich non-domain regions remain more laterally expanded and exhibit faster lipid diffusion. At the global level, the bacterial-inspired membrane displays greater bending rigidity than a fluid phosphatidylcholine reference, while ATR-FTIR measurements independently support increased lipid-chain ordering and denser membrane packing. Together, these findings establish that hopanoid-containing bacterial-inspired membranes can spontaneously generate coexisting physical environments whose dis- tinct local properties collectively determine global membrane behavior. The resulting experimental–computational framework provides a quantitative basis for investigating bacterial-like membrane heterogeneity and how local composition, dynamics, and mechanics govern responses to membrane-targeting perturbations.

## Introduction

The accelerating prevalence of antimicrobial resistance (AMR) has diminished the efficacy of many widely used antibiotics and has intensified the need for antibacterial strategies that are less vulnerable to classical resistance routes such as target mutation or drug inactiva- tion. The surveillance and synthesis efforts of the World Health Organization underscore the global scale of this challenge, including substantial resistance levels among common bacterial pathogens monitored internationally. ^1^ In parallel, there is renewed interest in therapeutically targeting conserved, indispensable cellular structures whose function emerges from collective physicochemical properties rather than a single stereospecific binding pocket. The bacterial cytoplasmic membrane is an attractive candidate in this regard because its essential roles in barrier function, organization of envelope proteins, and maintenance of electrochemical gradients depend on lipid packing, charge, and elasticity at the mesoscale. Accordingly, drug-discovery and medicinal-chemistry efforts have increasingly revisited membrane-active modalities-particularly those engaging bacterial anionic phospholipids as selectivity deter- minants, both as primary antibacterial mechanisms and as complements to conventional targets.^2,3^

Membrane lipids do not necessarily mix ideally: differences in headgroup chemistry, acylchain saturation, and molecular geometry can generate non-uniform lateral organization that segregates membranes into regions with distinct composition and physical state. In canon- ical model systems, such lateral segregation can yield coexisting liquid-ordered and liquid- disordered phases that differ in hydrocarbon chain order, packing density, viscosity, and lipid diffusion, with concomitant changes in bilayer thickness and mechanical response.^4^ Impor- tantly, these physical differences provide a plausible route by which membranes modulate biological function: by altering the energetic landscape for protein partitioning, oligomeriza- tion, and membrane-associated reaction kinetics and by shaping permeability and mechanical robustness. Experimental and theoretical work further emphasizes that the small size, short lifetimes, and compositional complexity of domains in living cells can obstruct direct charac- terization, motivating the use of controlled model membranes and quantitatively grounded approaches to connect composition, structure, and mechanics.^4^

Although bacterial membranes were long treated as comparatively homogeneous, an in- creasing body of work supports spatial heterogeneity and domain-like organization. One prominent and mechanistically interpretable example involves anionic phospholipids, es- pecially cardiolipin, whose enrichment at bacterial poles and septa sites has been widely reported and has been explicitly connected to curvature-dependent lipid sorting. In par- ticular, when bacterial membranes are forced into defined geometries like poles, cardiolipin microdomains preferentially localize to regions of high intrinsic negative curvature. This localization can coincide with the enrichment of specific membrane-associated proteins.^5^ Within this landscape, the concept of functional membrane microdomains has provided a useful framework by proposing that specific bacterial processes can be compartmentalized into membrane regions that locally concentrate proteins involved in signaling, transport and envelope-associated functions, while also explicitly acknowledging unresolved controversies over constituent lipids and organization principles.^6–8^ Together, these indicate that bacterial membrane heterogeneity can emerge in multiple forms and on different organizational levels, depending on the underlying lipid and protein determinants.

Most bacteria do not synthesize cholesterol, raising the question of how to achieve and tune membrane order and robustness under environmental stress and during envelope remod- eling. Another form of membrane heterogeneity may arise from hopanoids. Hopanoids, poly- cyclic isoprenoids produced by many bacterial lineages, have emerged as leading candidates for ”sterol analog” ordering lipids: in reconstituted membranes, hopanoids can promote the formation of ordered liquid phases in a manner that functionally converges with eukaryotic sterols, implying that polycyclic isoprenoids can stabilize a distinct, ordered fluid state capa- ble of supporting lateral heterogeneity.^9^ *In vivo* evidence provides a particularly compelling functional link: hopanoids were shown to determine the order of the bacterial membrane, and their depletion lowers the order of the membrane. In particular, it affects the efflux of multidrug-dependent energy.^10^ Beyond the immediate physiological implications, these findings motivate the hypothesis that hopanoid-dependent ordering can contribute to the formation and stabilization of laterally distinct membrane regions by coupling local compo- sition to emergent physical properties such as packing, thickness, diffusion, and mechanical resistance, precisely the parameters that set the energetic cost of protein insertion, confor- mational dynamics, and membrane-active compound binding.^10^

Progress in this area is limited by the scarcity of quantitatively validated bacterial mimicking membrane models that both reproduce compositional complexity and yield resolvable coexisting regions whose local structure and mechanics can be measured and compared.

This limitation is significant because membrane-active agents can alter the organization of the lipid domain as an early step of activity; high-speed atomic force microscopy, for ex- ample, has revealed that a small cationic antibacterial agent alters the organization and dynamics of lipid domains in *S.aureus* membranes, highlighting microdomains as physically and pharmacologically consequential structures.^11^

In this paper, we present an optimized bacterial-like membrane model that spontaneously forms distinguishable microdomains both *in vitro* and *in silico*. This enables an inte- grated characterization of domain formation using a combination of experimental studies and molecular dynamics (MD) simulations. Building on previous studies that demonstrated that hopanoids can induce ordered membrane states in simplified systems, ^12,13^ in our work, we extended this principle to a bacterial lipid mixture model inspired by lipid classes found in *Methylorubrum extorquens* membranes and provided a quantitative characterization of domain-like and non-domain regions in terms of packing, thickness, chain ordering, diffu- sion, and membrane flexibility. A central outcome of the study is the identification of dis- tinct regions with elevated hopanoid content. This supports the hypothesis that hopanoids may act as sterol analogs in bacteria and provides a reference point for analyzing how mi- crodomain mechanics and composition influence interactions with membrane-active antimi- crobial agents.

## Materials and methods

### Lipids and reagents

Phospholipids were obtained from Avanti Polar Lipids unless otherwise stated. These in- cluded POPC (1-palmitoyl-2-oleoyl-sn-glycero-3-phosphocholine), DPPG (1,2-dipalmitoyl-sn-glycero-3-phospho-(1’-rac-glycerol)), DiPE (1,2-dilinoleoyl-sn-glycero-3-phosphoethanolamine), and 18:1 CL (cardiolipin). Fluorescent lipid analogs 18:1-12:0 NBD-PC and TopFluor- cardiolipin were also purchased from Avanti Polar Lipids. The HOP (hopanoid, 22-hydroxyhopane) was purchased from Chiron. The DiIC_12_ membrane dye was purchased from Thermo Fisher Scientific. Atto488-DOPE was purchased from Sigma-Aldrich. Sucrose and HEPES (N- (2-hydroxyethyl)piperazine-N’-(2-ethanesulfonic acid)) were purchased from Sigma-Aldrich. Low-temperature agarose (melting temperature ≈ 65°*C*; gelling temperature ≈ 25°*C*) was purchased from Fisher Scientific. All aqueous solutions were prepared using ultrapure water produced by a water purification system (Millipore). The chemical structures of the lipids and probes used in this study are shown in Figures S1 and S2.

### Giant unilamellar vesicle electroformation

Giant unilamellar vesicles (GUVs) were prepared by electroformation in a glass chamber maintained at 45 °C.^14^ Lipids were dissolved in chloroform and mixed to yield a bacterial-like, *M. extorquens*-inspired working composition^15,16^ of 0.3PC:0.3PE:0.2PG:0.18HOP:0.02CL, where PC corresponds to POPC, PE to DiPE, PG to DPPG, CL to cardiolipin, and HOP to 22-hydroxyhopane. For deposition, the total lipid concentration was 1*mM* in chloroform. The membranes were fluorescently labeled by including 1mol % Atto488-DOPE and 1mol % DiIC. A 50*µl* aliquot of the 1*mM* lipid mixture was evenly distributed along the platinum electrodes. The solvent was evaporated and the residual chloroform was removed by drying the coated electrodes under vacuum for 3*h*. The electrodes were then assembled within the glass electroformation chamber and submerged in a non-conductive sucrose solution adjusted to 120 *mOsm*. An alternating electric field was applied using a 10 *Hz* square-wave AC waveform, starting at 1 *V* and then increasing by 1 *V* per hour until it reached 4 *V* .^17^ The field was maintained for a total of 24 *h*. Agarose immobilization was conducted according to Lira *et al*. protocol.^18^ Briefly, agarose was dissolved in 120 *mOsm* sucrose solution to obtain a concentration of 1% w/v. Vesicles and agarose were mixed equally in volume while agarose was still in the fluid state (around 35-40 °*C*).

### Vesicle fluctuation spectroscopy

Thermally induced shape fluctuations of GUVs can be used to determine mechanical prop- erties (bending rigidity and surface tension). The series of images of GUVs was recorded by camera-based fluorescence microscopy. A Leica TCS SPE (Leica, Wetzlar, Germany) was equipped with a 63×/1.30 ACS APO oil-immersion objective. Fluorophores were excited with an EL6000 illuminator (mercury metal halide bulb), and fluorescence emission was de- tected using I3 cut-off filter (BP 450-490 nm excitation filter, a 510 nm dichromatic mirror, and an LP 515 nm emission filter, transmitting light above 515 nm). Time-series movies were acquired at 1392x1040 pixels using a Leica DFC310 FX camera, typically comprising 3000 consecutive frames, with the imaging plane positioned at the vesicle center to maximize contour fidelity. The radii of the investigated vesicles ranged from ≈ 2 − 15*µm*. The samples were collected at a temperature of 22 ± 1 °*C*.

The bending rigidity of the membrane was obtained by fluctuation spectroscopy within the Helfrich elastic membrane framework. For each frame, the vesicle contour was identi- fied, and the instantaneous radial position of the membrane was extracted as a function of the polar angle. The contour extraction procedure was implemented using custom written C++/OpenCV software and was applied identically across all conditions. The resulting time-resolved radial fluctuations were processed in MATLAB using the average-based ap- proach. Here, angular autocorrelation functions were computed from the radial fluctuations and decomposed into a Legendre polynomial series; the mode-dependent fluctuation ampli- tudes were then fitted to obtain the bending rigidity. A detailed protocol has been described elsewhere.^19^

Three-dimensional z-stack images of GUVs were acquired with a Stellaris 8 confocal microscope (Leica, Wetzlar, Germany) using an HC PL APO 86×/1.20 water-immersion objective (Leica). Each image was recorded at 521 × 521 pixels with a pinhole size of 0.5 Airy units. Membranes were labeled with Atto488-DOPE and DiIC, and fluorescence excitation was provided by the white-light laser, with excitation wavelengths 488 nm for Atto488-DOPE and 549 nm for DiI. Emission signals were collected in separate detection channels with a hybrid (HyD) detector. Z-stacks spanning the full vesicle diameter were acquired for three-dimensional reconstruction and visualization of lateral membrane heterogeneity. All measurements were performed at 21 °C.

### Molecular dynamics simulations

Coarse-grained bilayer (CG) systems were constructed using CHARMM-GUI Martini Maker.^20^ Two membrane compositions were prepared: a PC-only bilayer (100% PC) and a bacterial- like model (BAC) containing 30% PC, 30% PE, 20% PG, 18% HOP, and 2% CL, which reflected the experimental setup.^15,16^ Each membrane system comprised 1000 lipids in to- tal. In this work, we employed the Martini 2.3P force field^21^ together with the polarizable water model. The bilayer systems were energy-minimized and equilibrated following the standard multi-step protocol. Production simulations were performed for 20 *µs* using a 20 *fs* integration time step. All simulations were run with GROMACS 2022.4.^22^ Before each production run, steepest-descent energy minimization was carried out. Simulations were per- formed in the isothermal–isobaric (NPT) ensemble. The temperature was controlled using the V-rescale thermostat at 295 *K* with a coupling time constant *τ_t_*= 1 *ps*. Pressure was con- trolled using the Parrinello–Rahman barostat^23^ applied semi-isotropically, with a coupling time constant of 12 *ps* and a compressibility of 3 ∗ 10*^−^*^4^ *bar^−^*^1^. Long-range electrostatics were treated using the reaction-field approach with a dielectric constant *ɛ_r_*=2.5. Each membrane simulation was repeated three times with random initial velocities to ensure reproducibility of the obtained results. Trajectories were processed using standard GROMACS tools and custom Python scripts based on the MDAnalysis library.

#### Area per lipid and membrane thickness

For each lipid, a reference point was defined at the midpoint of the selected headgroup beads (PO4,C1). The membrane plane (xy) was then tessellated using the Voronoi method to assign an instantaneous area to each lipid. From the collection of per-lipid areas over all frames, we constructed a histogram and fitted a Gaussian to determine the mean APL for the membrane. Membrane thickness was computed as the difference between the average headgroup height of the upper leaflet and that of the lower leaflet, and time averaged over the trajectory.

#### Bending rigidity

The bending rigidity *κ* was obtained by the real-space fluctuation (RSF) method.^24^ In this approach, the tilt angle of each lipid was defined as the angle between its director vector and the membrane normal. The director vector was drawn from the headgroup midpoint to the tail midpoint (where the tail midpoint is the average position of the terminal tail beads). The lipid splay was characterized by the relative orientation of the neighboring lipid directors. The tilt and splay angles distributions were collected on all lipids and frames, and the *κ* modulus was extracted by fitting these distributions according to the RSF formalism. ^24^

#### Order-parameter and lipid enrichment maps

Lipid tail order parameters were calculated using the *gorder* Python API.^25^ Briefly, for each lipid tail segment, the angle *θ* between the segment (bond) vector and the membrane normal was used to compute *S* = ^1^⟨3 cos^2^ *θ* − 1⟩. These per-bond order values were averaged to give a per-lipid order parameter and then averaged over time and lipids to yield the global order. To visualize spatial heterogeneity, order-parameter maps were generated on a (x, y) grid; the order from each lipid bond was averaged and projected on the membrane grid.

### Fluorescence recovery after photobleaching

The GUVs were prepared for Fluorescence recovery after photobleaching (FRAP) by elec- troformation as stated and dyed with NBD-PC. For imaging, vesicles were immobilized in ≈ 1% agarose to limit lipid mobility. The experiments were carried out on a Leica SP8 confocal microscope with an oil-immersion 64×/1.20 W objective at room temperature. The white-light laser was tuned to 488 nm for excitation. A pre-bleach sequence of 10 frames and a post-bleach sequence of 100 frames were recorded at low laser power (3%). Bleaching was executed as a single high-intensity (90%) 488 nm pulse focused on a circular ROI at the equatorial plane. The normalized recovery curve was fitted using the analytical Soumpasis solution to obtain the mobile fraction and the diffusion coefficient.^26^ The analysis was carried out with custom Python scripts.

### Attenuated total reflectance Fourier transform infrared spectroscopy

Spectroscopic measurements were conducted using attenuated total reflectance Fourier trans- form infrared spectroscopy (ATR-FTIR) on a Nicolet 6700 FTIR spectrometer (Thermo Sci- entific, USA) fitted with an ATR accessory and a temperature-regulated diamond top-plate from PIKE Technologies. The analyzes were performed under continuous dry air purging. Sample preparation involved depositing 10 *µl* of GUVs suspension onto the diamond crys- tal surface and allowing complete solvent evaporation before spectral acquisition using the dry-film methodology. Infrared spectra were recorded over the 3600–400 *cm^−^*^1^ range by co-adding 128 interferograms at 4 *cm^−^*^1^ spectral resolution (Figure S3). Each sample was measured four or more times, and the averaged spectrum was used for subsequent analy- sis. All measurements were carried out at a controlled temperature of 22 °*C*. Background spectra of the clean diamond crystal in air were collected before each measurement (512 scans, 4 *cm^−^*^1^ resolution). To optimize signal-to-noise ratios, samples were prepared with lipid concentration of 1 *µM* . Data processing was performed using OriginPro 2024b software (OriginLab Corporation, Northampton, Massachusetts, United States), incorporating auto- matic atmospheric compensation, baseline correction, and spectral normalization relative to the *CH*_2_ stretching band at 1450 *cm^−^*^1^. Second-derivative spectra were smoothed using a Savitzky-Golay filter with a polynomial order of 15.^27^

### Statistical analysis

Statistical analysis was performed in Python using the statsmodels library.^28^ All experimen- tal measurements were repeated at least three times, and the MD system included three independent replicas. Results are presented as mean values with the corresponding stan- dard error. Statistical significance was assessed by one-way analysis of variance (ANOVA), followed by Tukey’s post hoc test for multiple comparisons. Differences were considered statistically significant at p *<* 0.05.

## Results and discussion

### Characterization of GUV domain morphology

Confocal imaging of GUVs prepared from the bacterial-mimetic mixture (BAC) revealed lateral heterogeneity that was resolved on the micrometer scale (Fig. 1 A,B). In three- dimensional reconstructions of individual vesicles, membrane fluorescence did not appear spatially uniform; instead, two-color labeling (Atto488-DOPE in the red channel and a DiL dye in the green channel) reported extended membrane regions with distinct intensity pat- terns. The observed contrast was stable across the acquisition intervals, thus, it reflects persistent lateral reorganization rather than transient imaging noise or fluctuations. Impor- tantly, the resulting patterns were rather not punctate or weakly heterogeneous, with mostly one or two continuous membrane regions per vesicle. This behavior is consistent with stable domain-like lateral segregation in the membrane system rather than a highly fragmented pat- tern (Fig. 1 C). The present microscopy data are consistent with stable lateral segregation, although they do not by themselves assign the coexisting regions to a specific thermodynamic phase state. For this reason, we refer to them throughout the manuscript as domain-like and non-domain regions. The mean sizes of the vesicles obtained equaled 6.87 ± 0.9*µm* in diameter (see Fig. S3). The presence of resolvable domain-like regions in a free-standing bacterial-inspired bilayer is notable, since many proposed bacterial membrane microdomains *in vivo* remain difficult to verify directly due to the complexity of their composition, small size, or methodological limitations. A recent review of bacterial membrane biophysics iden- tified lateral microdomains as a new organizational principle and discussed, in particular, isoprenoid lipids as potential contributors to membrane organization, while also emphasizing that the mechanistic principles of this process remain incompletely defined. ^7^ In this context, the imaging results shown in Figs. 1 A and 1 B indicate that the present lipid mixture, which contains a hopanoid component, is sufficient to induce macroscopic lateral segregation in the absence of proteins or active cellular processes, providing an easily studied reference system for further analysis of the mechanisms involved.

**Figure 1:**
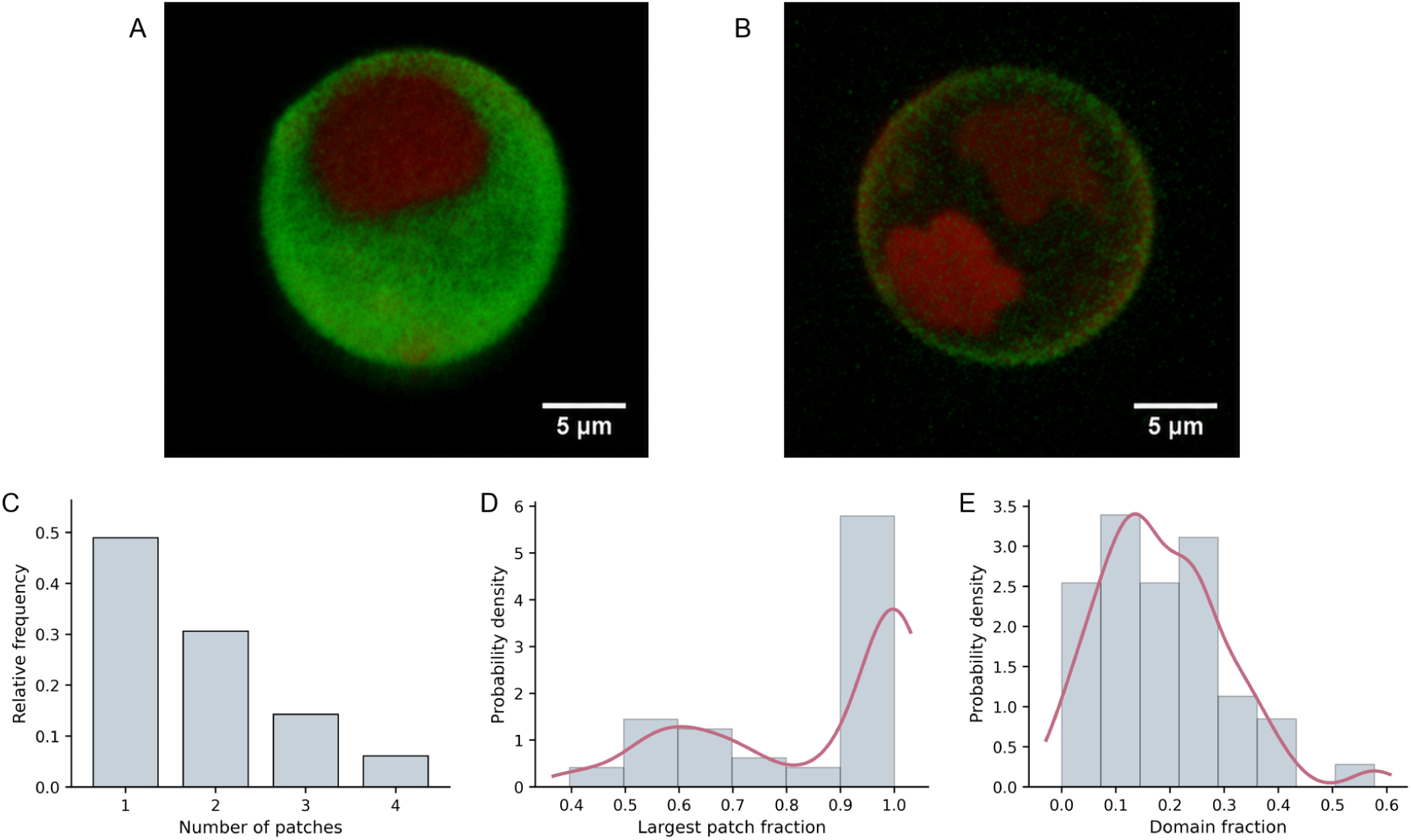
Microdomain formation in the bacterial-mimetic membrane. (A,B) Representative confocal images of GUVs showing lateral segregation of the fluorescent probes and the pres- ence of distinguishable membrane microdomains. (C) Distribution of the number of patches per vesicle. (D) Distribution of the largest patch fraction, defined as the fraction of total domain area occupied by the largest domain within an individual vesicle. (E) Distribution of the domain fraction in the membrane band, describing the fraction of vesicle membrane assigned to the domain phase.

To complement qualitative inspection, we quantified domain morphology using connected- component analysis of the segmented domains derived from the confocal signal (Fig. 1 C-D). The distribution of the number of patches per vesicle was dominated by vesicles containing a single large patch (Fig. 1 C). Several samples exhibited two patches, whereas vesicles with three or four patches were relatively infrequent. Overall, this distribution supports the practical observation that most vesicles display “one” or “two-domain” phenotypes rather than extensive fragmentation.

To capture how the domain is partitioned among patches within each vesicle, we analyzed the largest patch fraction, defined here as the fraction of the total domain-assigned area (the total segmented domain signal within the analyzed membrane band) contributed by the single largest connected patch within a given vesicle. The resulting probability density shows a strong enrichment near unity (Fig. 1D), indicating that in many vesicles, most of the domain is consolidated into a single dominant patch. The same distribution exhibits a secondary elevation around 0.5–0.7, consistent with a subset of vesicles in which two patches contribute comparably to the total domain area. In practical terms, the combination of a high near-1 peak with a shoulder near 0.5 is the expected signature of a population dominated by one large domain, along with a secondary population in which two domains of similar size persist.

Next, we quantified the overall abundance of domains using the domain fraction, defined as the fraction of the analyzed membrane band classified as domain (Fig. 1 E). The mean domain occupancy equals 18%±1.5% of membrane band. The distribution is skewed towards relatively low fractions, with most vesicles falling in the approximate range of ≈ 0.05–0.30 and a tail extending to higher values (approaching ≈ 0.4). This is consistent with the ex- pectation that electroformation can yield modest vesicle-to-vesicle heterogeneity in effective composition and tension, both of which shift the balance between coexisting membrane en- vironments. The observed predominance of one or two large patches is also consistent with the generic coarsening behavior in phase-separating GUV membranes. Once two fluid phases coexist in a free-standing bilayer, the domains typically grow through a combination of collision–coalescence and (depending on the conditions) evaporation–condensation, reducing the total interfacial length as the system evolves.^29^ The present patch statistics therefore sup- port an interpretation in which the GUV microdomains represent a later-stage, coarsened morphology rather than an early, highly nucleated pattern.

### Domain segregation and local ordering in MD simulations

The experimental observations are mirrored by coarse-grained MD simulations performed on a compositionally matched bilayer. In the upper-leaflet compositional snapshot (Fig. 2 A), the membrane resolves into PE-rich, non-domain environment and a HOP/PC/PG/CL- enriched, domain-like environment. The PE enrichment map (Fig. 2 B) independently supports the same partitioning by explicitly highlighting DIPE-rich and depleted regions. Together, these indicate that the simulated membrane does not remain laterally mixed but instead undergoes composition-dependent segregation into two coexisting membrane envi- ronments. Beyond composition, the simulated membrane also exhibits spatial heterogeneity in lipid ordering. An order-parameter map (Fig. 2 C) shows adjacent regions with higher order contrasted against a background with lower order. Although the order metric is a coarse-grained analogue rather than a direct *CH* bond order parameter, its spatial pattern aligns with the compositional classification that the regions corresponding to the domain- like environment present a higher order than the PE-enriched non-domain one (Fig. 2 A-C). The internal consistency across compositional and order representations is important as it indicates that domain identification does not arise solely from an arbitrary encoding of lipid identity but rather that the segregation is accompanied by a physically interpretable difference in local chain organization. This coupling between the HOP-enriched composition and the increased order is aligned with prior work showing that hopanoids can behave as sterol-like order modulators and can promote liquid–liquid phase separation in model membranes. In previous work there was reported that hopanoids can form liquid-ordered phases and promote liquid–liquid phase separation in membranes, providing a mechanis- tic rationale for hopanoid-associated lateral segregation.^10,13^ Independent experiments on diplopterol-containing membranes further reported that a hopanoid can induce liquid–liquid phase segregation while increasing lipid compaction, consistent with an ordering role rather than a simple rigidification into a gel-like state.^12^ In this context, the present simulations provide a direct, composition-resolved view of how a PE-rich region can coexist with a HOP- containing region within the same bilayer.

**Figure 2:**
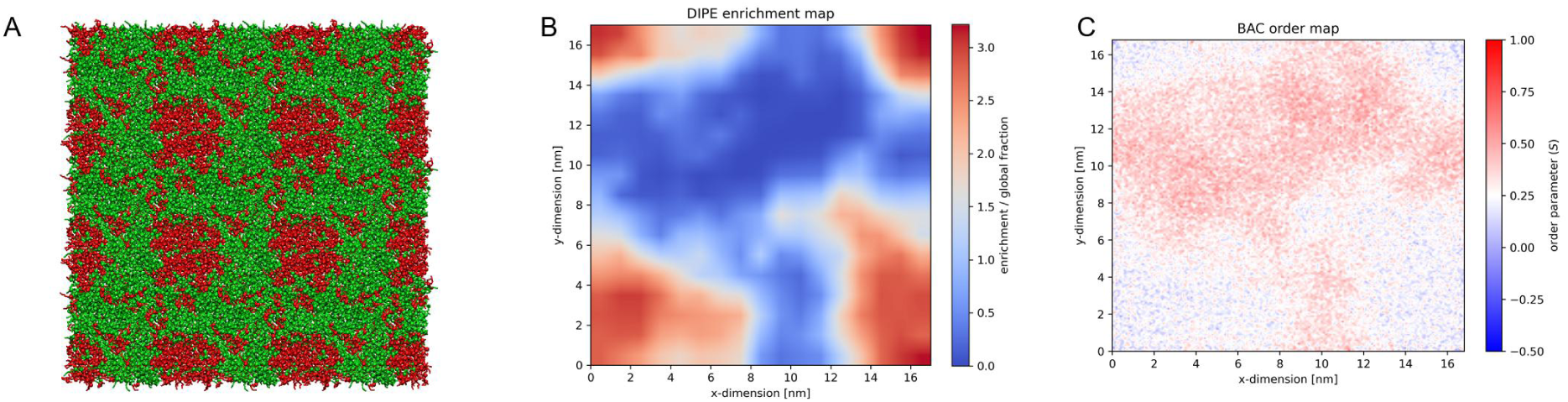
MD visualization of lateral heterogeneity in the bacterial-mimetic membrane. (A) Representative MD snapshot, with PC/PG/HOP/CL beads shown in green and PE beads in red. (B) PE enrichment map showing DIPE-rich and DIPE-poor membrane regions. (C) Order-parameter map showing local differences in lipid ordering that coincide with the segregated membrane environments.

Taken together, hopanoid-containing bacterial-mimetic bilayers can reproducibly exhibit lateral segregation that is detectable both experimentally and computationally, and the sim- ulated membrane regions classified by composition correspond to a physically meaningful separation in molecular order. The applied approach provides a rigorous starting point for subsequent analysis in which dynamics and mechanics can be quantified and related to hy- potheses about hopanoid-driven ordering^12,30^ and its potential relevance to the organization of bacterial membranes.

### Mobility and diffusion

To determine whether the domain-forming state identified in Fig. 1, 2 is accompanied by altered membrane dynamics, we next compared lipid mobility in the BAC and in the PC control using FRAP and MD simulations (Fig. 3 and Fig S4). PC membrane was used as a reference system since such bilayers are widely employed as experimentally tractable and physically well-characterized benchmarks in GUV biophysics. The PC diffusion obtained followed what was previously reported in the literature.^31,32^ FRAP showed higher apparent mobility in the BAC membrane than in the PC control (Fig. 2 A). This result indicates that the mixed BAC membrane does not behave as a uniformly condensed bilayer despite the presence of laterally segregated regions. Such behavior is consistent with the general view that bacterial membranes can maintain substantial lateral dynamics even in the presence of compositional heterogeneity and domain-like organization.^7^

**Figure 3:**
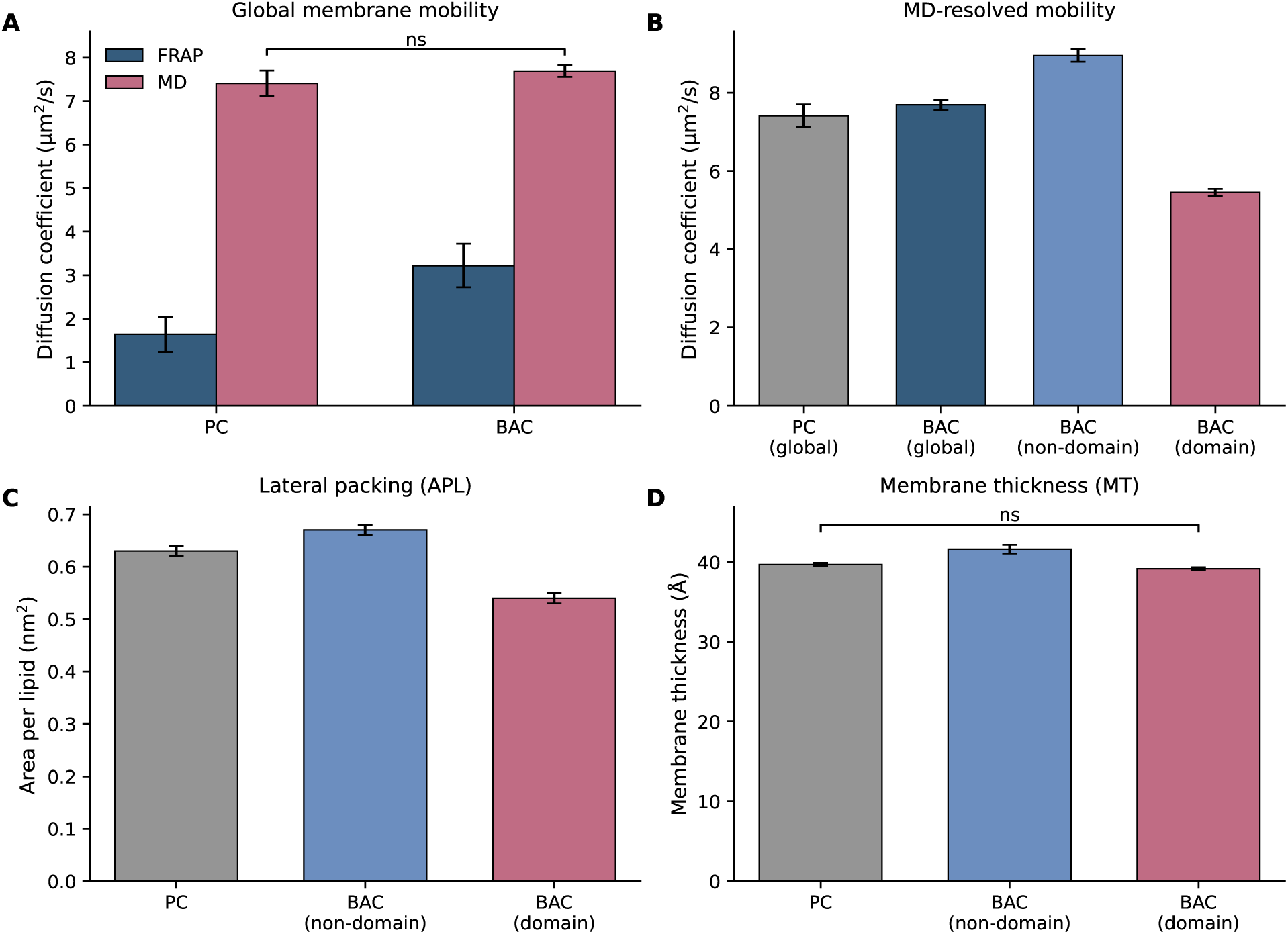
Membrane mobility and structural parameters of the PC reference and bacterial- mimetic membrane. (A) Global membrane mobility determined by FRAP and MD simula- tions for the PC and BAC systems. (B) MD-resolved lateral diffusion coefficients for the PC and BAC membrane represented as global, non-domain, and domain environments. (C) Area per lipid for the PC and BAC non-domain and domain regions. (D) Membrane thickness for the PC membrane and BAC non-domain and domain regions. Non-significant comparisons (p *>* 0.05) are indicated as *ns*.

The MD analysis refined this observation by showing that the BAC membrane is not dynamically uniform. When diffusion was averaged over the whole membrane, the difference relative to PC was small (Fig. 3 A,B). FRAP reports an effective diffusion that is weighted by the probe’s partitioning and by the connectivity of the diffusing phase, whereas MD provides access to region-specific diffusion within coexisting membrane environments. The nondomain environment showed the highest lateral diffusion coefficient, while the domain-like was markedly less mobile, with the PC control occupying an intermediate position (Fig. 3 B). The main result of the simulation is therefore not a large shift in global membrane diffusion but the emergence of dynamic heterogeneity within the membrane plane. This regional difference in mobility was accompanied by a pronounced difference in APL. As shown in Fig. 3 C, the DIPE-rich environment had the largest APL, the domain phase the smallest, and PC again an intermediate value. This matched the ordering of diffusion; the most expanded membrane environment was also the most mobile, whereas the most condensed environment was the least mobile. Here, APL provides the clearest structural correlate of lateral diffusion. It stays in line with membrane biophysics more broadly, where lateral diffusion is commonly associated with free area, lateral packing, and molecular ordering within the bilayer.^33^ Moreover, in laterally heterogeneous membranes, the relevant dynamic unit is often the local phase or domain rather than the membrane average.^34^

It should be noted that hopanoids have been described as sterol surrogates in bacteria and have been shown to promote membrane ordering in a sterol-like manner.^9,10,35^ As previously stated, diplopterol has been reported to induce phase segregation, increase lipid compaction, and reduce membrane permeability while retaining membrane fluidity and compressibil- ity.^12,13^ Our data are consistent with that framework. The slower domain phase in the bacterial-mimetic membrane is also the more laterally condensed phase, whereas the DIPE- rich environment remains more expanded and mobile. Thus, the coexistence of fast and slow membrane regions in our simulations is compatible with the expected effect of hopanoid- containing mixtures, namely the formation of locally differentiated membrane environments (Fig. 3 B). MT followed the same qualitative ordering and further supported structural differentiation between the two membrane environments, with MT = 41.62 ± 0.57Å in non- domain regions, MT = 39.69 ± 0.21Å in PC, and MT = 39.15 ± 0.28 Å in the domain regions (Fig. 3 D). Although MT therefore provides an independent structural signature of the two phases, the strongest mechanistic association with mobility in the present dataset is observed for APL: the largest separation between environments is in lateral packing (0.67 vs 0.54 *nm*^2^), which co-varies with the clearest separation in diffusion (8.95 vs 5.45 *µs*^2^ ∗ *m^−^*^1^). The thickness differences are smaller in magnitude, but remain consistent with two coexisting membrane states rather than a single laterally uniform bilayer.

Importantly, the PC control values obtained in this study were consistent with previously reported ranges, including FRAP mobility,^32^ lateral diffusion,^31^ APL, and MT,^3,31^ support- ing the validity of the experimental and computational baselines used here. Notably, the diffusion values obtained for the bacterial-mimetic membrane also fall within ranges reported previously for bacteria, suggesting that this model reproduces selected dynamic features of biological membranes in a simplified and experimentally tractable setting. Recent measure- ments in *Bacillus subtilis* and *Staphylococcus aureus* reported diffusion coefficients^36^ com- parable to the values observed here for the bacterial-like membrane, whereas FRAP-based studies in *Escherichia coli* have reported diffusion coefficients spanning approximately 3–14 *µ*m^2^/s, depending on the system, probe, and experimental conditions.^37^ Although these liter- ature values do not refer specifically to bacterial-mimetic model membrane however, provide a useful biological context for interpreting the present data and suggest that the mobility regime captured by our system may offer insight into the dynamic behavior of bacterial membranes. While the present model does not recapitulate the full complexity of a living bacterial envelope, its diffusion behavior remains within a biologically plausible range and supports its use as a reduced platform for studying membrane heterogeneity and domain-associated effects. In this context, the fluorescent lipid analogs used here are interpreted as reporters of membrane organization, and the fluorescence-based observations are considered together with the orthogonal evidence provided by molecular simulations, ATR-FTIR, and membrane-mechanical measurements.

Together, the results are consistent with the growing view that the organization of the bacterial membrane should be understood in terms of physically distinct lateral environments that differ not only in composition but also in mobility and local material properties.

### Mechanical description of systems

To further evaluate whether microdomain formation in the bacterial-mimetic membrane is accompanied by changes in membrane mechanics, we next compared bending with vesicle fluctuation spectroscopy (Fig. 4). The PC control bending rigidity obtained here falls within the literature range for such membranes by flickering/fluctuation-based approaches^14,38^ and simulation-based fluctuation analyzes.^24,39,40^

**Figure 4:**
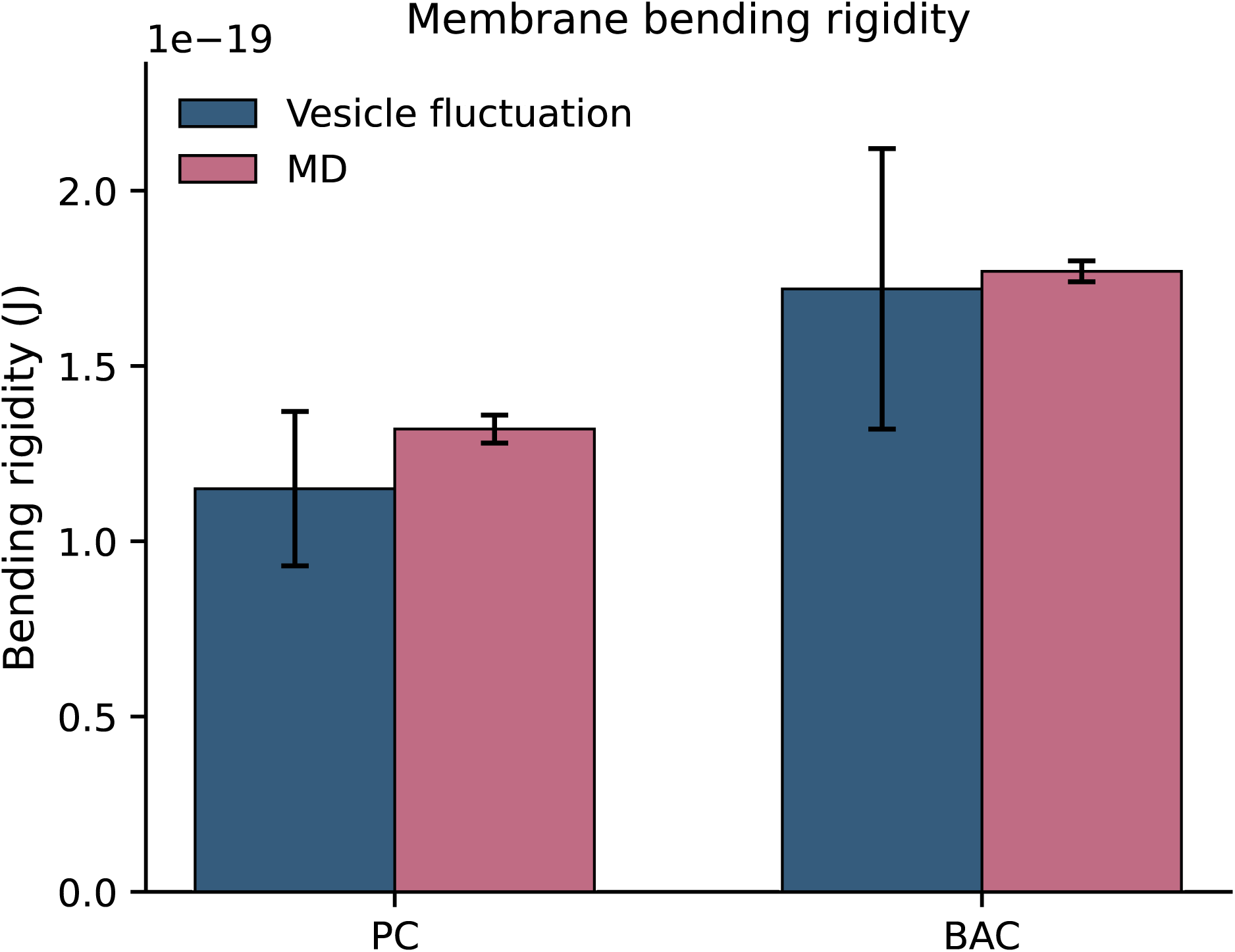
Bending rigidity measured by vesicle fluctuation analysis and estimated from MD simulations for the PC reference and BAC membrane. All pairwise comparisons between PC and BAC were statistically significant (p*<* 0.05)

The bacterial-mimetic membrane was consistently stiffer than the PC reference in both approaches. The GUV bending modulus increased from 1.1 ∗ 10*^−^*^19^*J* for the PC control to 1.7 ∗ 10*^−^*^19^*J* for the BAC membrane, while in the corresponding MD analysis 1.3 ∗ 10*^−^*^19^*J* and 1.8 ∗ 10*^−^*^19^*J* (Fig. 4). This agreement is notable because bending rigidity is a global mechanical descriptor of the bilayer. The fact that the same ordering was recovered by vesicle fluctuation analysis and MD indicates that the higher stiffness of the BAC membrane is an intrinsic property of the membrane state established in this system. Changes in lipid composition and lateral organization are well known to alter membrane bending elasticity, and increased membrane order is commonly associated with higher resistance to bending. It should be noted that the BAC membrane acts as a mechanically reinforced relative to a canonical fluid PC bilayer, while remaining within a regime typical of biologically relevant soft membranes rather than gel-like solids.^38^

The mechanical contrast between BAC and PC gains mechanistic meaning when interpreted alongside the mobility and packing results described in the prevous subsections. FRAP showed that the BAC membrane exhibits high apparent tracer mobility relative to PC, whereas MD did not indicate a large shift in a single global-average diffusion coefficient. Instead, it resolved the BAC bilayer into dynamically distinct environments. PE-rich re- gions were associated with faster diffusion and larger APL, while domain-like regions were associated with slower diffusion and smaller APL. This pattern argues against a single homo- geneous physical state and supports a model in which the BAC membrane comprises at least two coexisting membrane environments, a more mobile, laterally expanded DIPE-rich envi- ronment and a more condensed, slower environment enriched in the remaining components.^41^

In this context, a higher global bending rigidity should not be interpreted as a statement that “the entire membrane is locally rigid everywhere”. Rather, the experimentally mea- sured *κ* is an effective, system-level resistance to bending that integrates contributions from coexisting regions and their coupling across domain boundaries. A membrane can therefore remain highly mobile in a percolating, fluid environment (as sensed by FRAP, depending on probe partitioning and connectivity) while still exhibiting increased resistance to bending because a substantial fraction of the surface is occupied by a more condensed environment with higher local order and packing density. This qualitative reconciliation is consistent with general structure–mechanics relationships in lipid bilayers, where tighter packing and increased effective thickness tend to increase bending rigidity, while increased unsaturation and chain disorder tend to soften membranes.^42,43^

Additionally, results from ATR FTIR spectroscopy also suggest that the BAC membrane differs from PC in its hydrocarbon-chain organization (Figs. S5-S8; Table S1). Examination of the ATR-FTIR spectra in the 2980-2820 *cm^−^*^1^ region, corresponding to the symmetric and asymmetric stretching vibrations of the *CH*_2_ and *CH*_3_ groups, revealed systematic differences between the PC and BAC membranes (see Figure S6 and Table S1). The BAC membrane exhibited consistent shifts toward lower wavenumbers for all methylene and methyl stretching modes relative to PC. Specifically, the asymmetric *CH*_2_ stretching band was observed at 2920 *cm^−^*^1^ in PC and shifted to 2917 *cm^−^*^1^ in BAC, while the symmetric *CH*_2_ stretching mode decreased from 2851 *cm^−^*^1^ to 2849 *cm^−^*^1^. These spectroscopic shifts are indicative of increased conformational order and enhanced all-trans rotamer populations within the BAC membrane acyl chains, suggesting a more rigid and densely packed lipid arrangement.^44–46^ Further analysis of the *CH*_2_*/CH*_3_ deformation region (1500-1400 *cm^−^*^1^) provided additional evidence supporting these observations (see Figure S7 and Table S1). The *CH*_2_ scissoring mode at 1468 *cm^−^*^1^ in PC shifted to 1466 *cm^−^*^1^ in BAC, accompanied by a corresponding displacement of the shoulder band from 1456 *cm^−^*^1^ to 1455 *cm^−^*^1^. Additionally, examination of the *CH*_2_ rocking vibration in the 750-650 *cm^−^*^1^ region (see Figure S8) revealed a shift from 722 *cm^−^*^1^ (PC) to 721 *cm^−^*^1^ (BAC), providing complementary evidence of increased chain ordering in the BAC membrane. The lower frequency positions characteristic of BAC are associated with increased populations of all-trans conformations and reduced membrane fluidity.^47^

Importantly, these FTIR spectra should be interpreted as reflecting the conformational order and packing (structural properties averaged over the entire complex), rather than as a direct measure of lateral diffusion across the entire membrane. FTIR can sensitively report changes in the acyl-chain conformational state, and in the case of mixtures, it enables the identification of coexistence regimes and the distinct melting behavior of individual com- ponents.^44,48^ However, it does not allow to determine whether the overall lateral mobility detected by the fluorescent probe in the FRAP has decreased or increased. In the current model, the most reasonable interpretation is that the BAC membrane exhibits, on average, increased conformational order because it contains a region of higher density (as also seen in the MD order map - see Figure 2 C), while simultaneously exhibiting elevated apparent tracer mobility at the vesicle scale due to spatial heterogeneity and weighting of FRAP by probe partitioning and phase connectivity. In conclusion, within that framework, the ele- vated bending rigidity of the BAC membrane can be understood as a macroscopic mechanical consequence of the introduction of a more condensed, hopanoid-containing environment that coexists with a more laterally expanded, PE-rich environment. The key implication is not simply that the BAC membrane is stiffer than PC, but that stiffness, mobility, and chain or- der collectively indicate a laterally heterogeneous membrane whose global mechanics emerge from the coexistence of environments with distinct packing and dynamical properties.^9,49,50^

## Conclusions

Here, we established an optimized hopanoid-containing bacterial-like membrane model that spontaneously forms distinguishable domain-like regions both *in vitro* and *in silico*. By combining confocal imaging of free-standing GUVs with coarse-grained molecular dynamics simu- lations, we obtained a consistent description of lateral membrane heterogeneity across exper- imental and computational scales. The model extends previous observations that hopanoids can promote ordered membrane states in simplified systems to a more complex bacterial-like lipid mixture^12,13^ and provides a tractable platform for examining how membrane composi- tion gives rise to domain formation, altered mobility, and changes in membrane mechanics. Several conclusions follow from the present study. First, the BAC membrane reproducibly formed extended membrane domains in GUVs. These domains were not observed as a highly fragmented population of small patches, but rather as one dominant domain or, less frequently, two large domains within a vesicle. This pattern indicates that the mem- brane mixture does not remain laterally uniform under the examined conditions and instead undergoes robust segregation into coexisting membrane regions. Second, the same compo- sitionally matched system displayed lateral segregation in MD simulations, where PE-rich, non-domain regions and PC/PG/HOP/CL-rich domain-like environments emerged within the same bilayer. The agreement between microscopy and simulation indicates that the ob- served heterogeneity reflects an intrinsic property of the membrane mixture rather than a feature specific to one method. In this sense, the present model provides an experimentally accessible reference system for studying bacterial membrane microdomains, whose direct characterization *in vivo* remains challenging.^7^

Third, the segregated regions were physically distinct. In the simulations, the domain-like environment exhibited higher local order, lower lateral diffusion, and smaller area per lipid, whereas the PE-rich environment remained more mobile and more laterally expanded. These results support a model in which the BAC membrane comprises at least two coexisting membrane environments with different packing and dynamic properties. Importantly, the diffusion values obtained for the bacterial-mimetic membrane remained within the range re- ported for bacterial membranes,^36,37^ which provides an additional indication that the model captures biologically plausible dynamic behavior rather than only reproducing generic phase separation in an artificial lipid mixture. In this sense, diffusion serves not only as a descriptor of membrane heterogeneity, but also as an independent validator of the bacterial-like char- acter of the system. More generally, the observed association between mobility and lateral packing is consistent with the established dependence of membrane diffusion on free area and local membrane environment.^33,34^

Fourth, the BAC membrane was mechanically stiffer than the PC reference. This trend was recovered independently by vesicle fluctuation spectroscopy and MD-based bending analysis, providing consistent evidence that the BAC membrane has a higher resistance to bending. ATR-FTIR measurements supported this interpretation by showing shifts in the hydrocarbon-chain vibrational region consistent with increased conformational order and denser chain packing in BAC relative to PC.^44–47^ Taken together, these observations indi- cate that domain formation in the BAC membrane is associated not only with compositional segregation, but also with measurable differences in order, mobility, and mechanics. This interpretation is further supported by the broader relationship between membrane packing and bending elasticity described for lipid bilayers.^42,43^

A central outcome of the study is the identification of distinct membrane regions associated with elevated hopanoid content and increased local order. This supports the hypothesis that hopanoids can act as sterol analogs in bacteria and suggests that their role extends be- yond general membrane stabilization to the generation of laterally differentiated membrane states.^9,10,12,13^ Although the present model does not resolve the full structural complexity of native bacterial membranes, it captures a physically meaningful hopanoid-containing mem- brane state in a controlled and experimentally accessible setting. More broadly, the present model establishes an experimentally tractable and computationally resolved platform for linking bacterial-like membrane heterogeneity to membrane function. Given growing evi- dence that hopanoid-containing membranes generate laterally distinct physical environments and that bacterial membrane domains can shape the action of membrane-active antimicro- bials, this system provides a useful basis for mechanistic studies of how local composition, packing, and mechanics control membrane susceptibility to perturbation.^7,9,49,50^

## Acknowledgement

M.R. thanks to the National Science Centre, Poland (grant no. 2022/45/N/NZ9/02130) for financial support. The authors express their sincere gratitude for the valuable discussions and insights provided by Dr. Micha-l Majkowski from the University of Wroclaw.

## Data Availability

MD simulations and trajectory analyses were performed using commonly used programs, i.e. GROMACS or VMD. Analyses were supported and performed with Python and Matlab scripting. All procedures employed in this work are described in the Methods section. All system information, input parameter files and experimental measurements, will be made publicly available upon publication in a public repository and the Supporting Information.

## Author Contributions

MR conceived the study and designed the research. MR, MK, DD developed the experimen- tal membrane model and performed the GUV imaging, FRAP, ATR-FTIR, and membrane fluctuation measurements. MR performed the molecular dynamics simulations and compu- tational analyses. MR, NS analyzed the experimental and simulation data and prepared the figures. MR supervised the project and contributed to the interpretation of the results. MR wrote the original draft of the manuscript. All authors reviewed and edited the manuscript, discussed the results, and approved the final version.

## Supporting Information Available

MD simulations and trajectory analyses were performed using commonly used programs, i.e. GROMACS or VMD. Analyses were supported and performed with Python and Matlab scripting. All procedures employed in this work are described in the Methods section. All system information, input parameter files and experimental measurements will be made publicly available upon publication in a public repository and the Supporting Information.

