## Supplementary material for "Bacterial-Mimetic Membranes Reveal Hopanoid-Associated Domain-Like Lateral Heterogeneity": Suplemental File

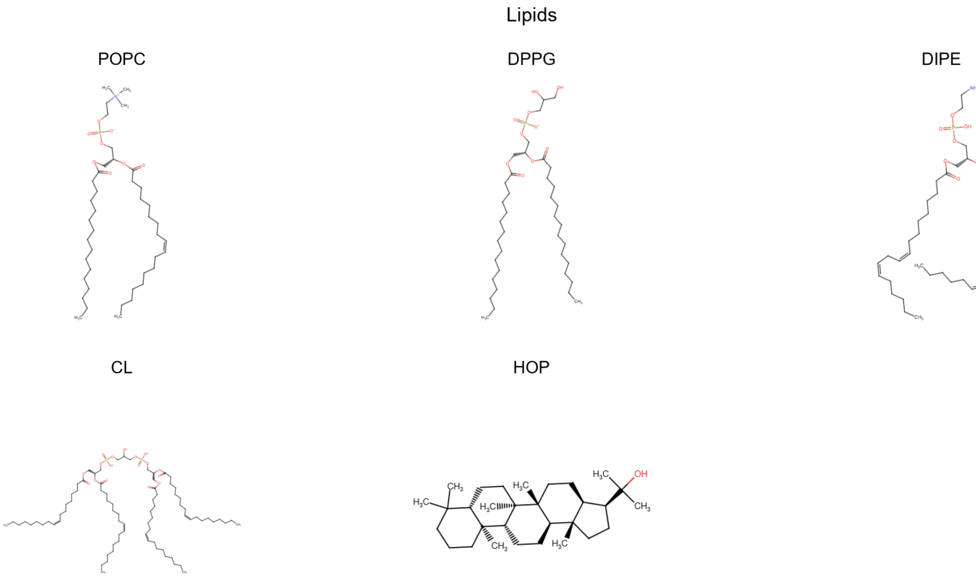


Figure S1: Chemical structures of the lipids used in this study.

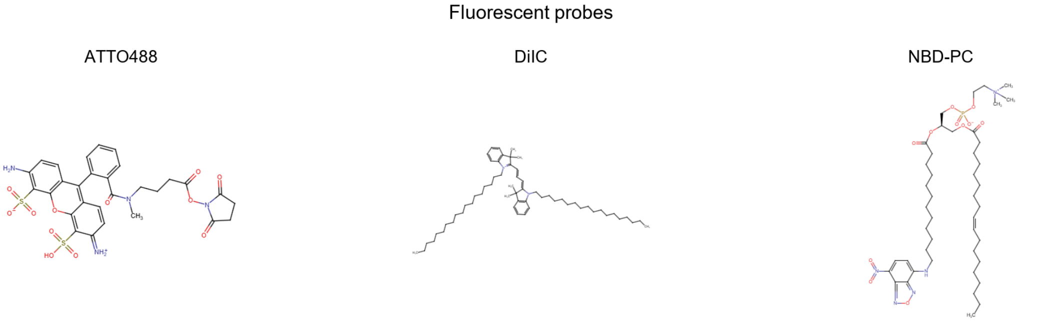


Figure S2: Chemical structures of the fluorescent probes used in this study.

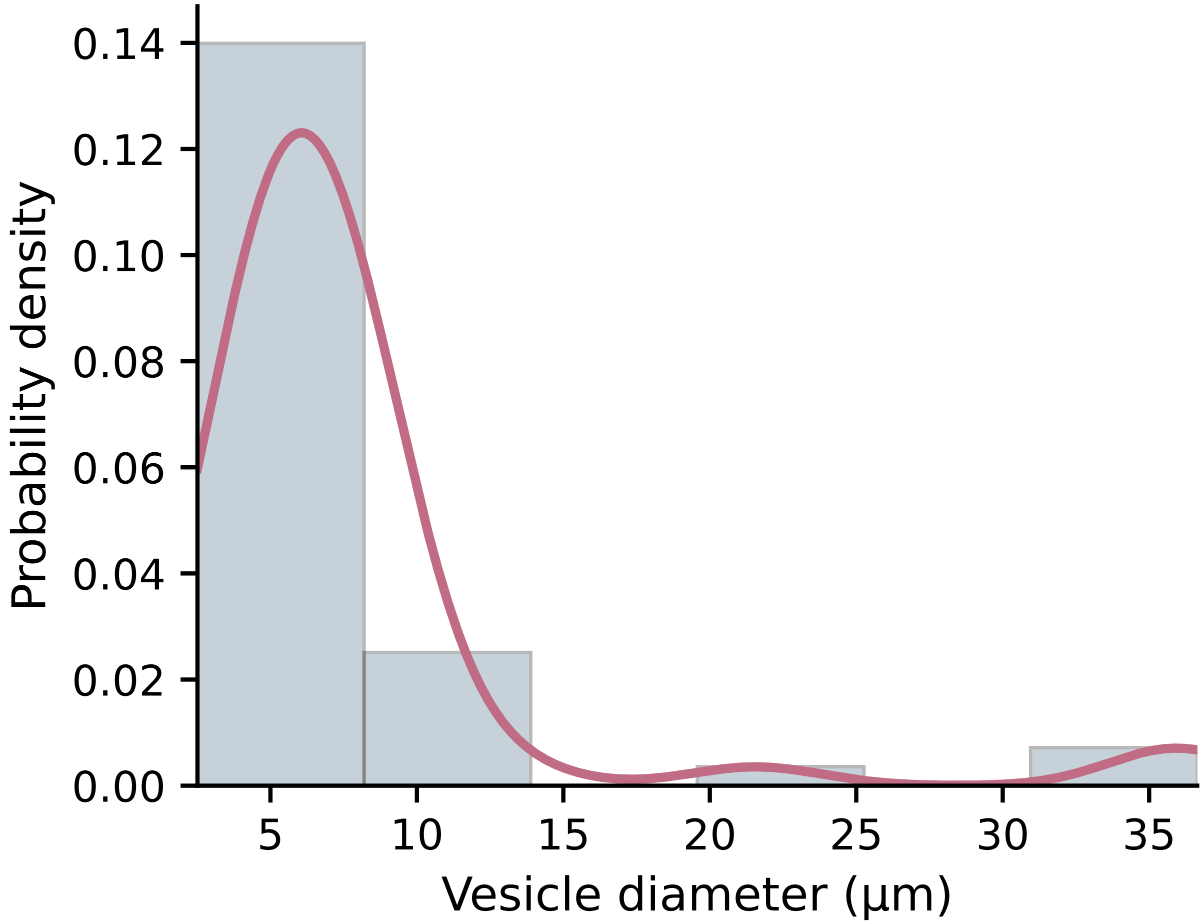


Figure S3: Distribution of vesicle diameters determined from microscopy images. The histogram shows the normalized probability density, while the solid line represents the kernel density estimate of the diameter distribution.

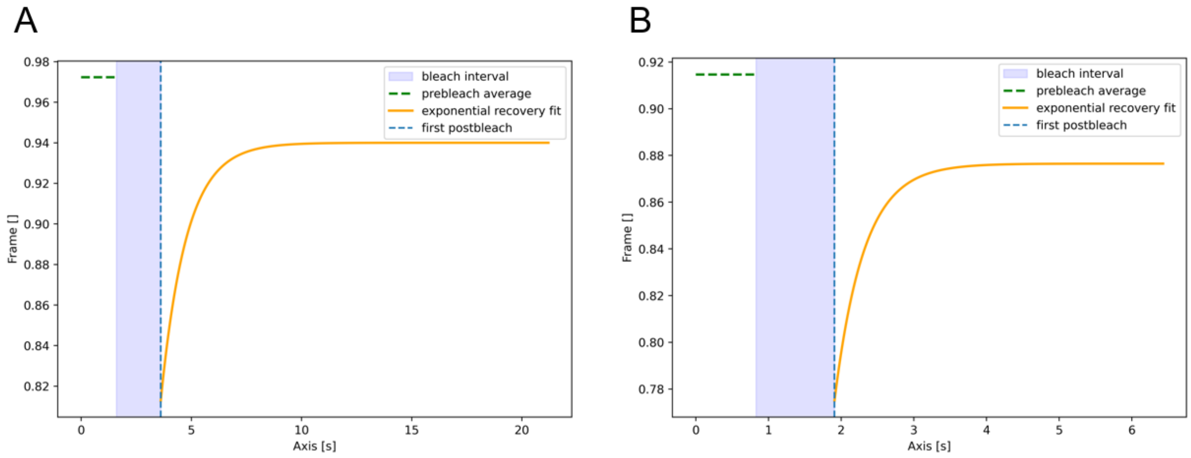


Figure S4: Representative FRAP recovery curves for the bacterial-mimetic and PC control membranes. (A) Representative fluorescence recovery curve for the BAC membrane. (B) Representative fluorescence recovery curve for the PC membrane. The shaded region indicates the bleaching interval, the dashed horizontal line marks the prebleach average, the blue dashed line marks the first postbleach time point, and the orange curve shows the exponential recovery fit used for the extraction of diffusion parameters.

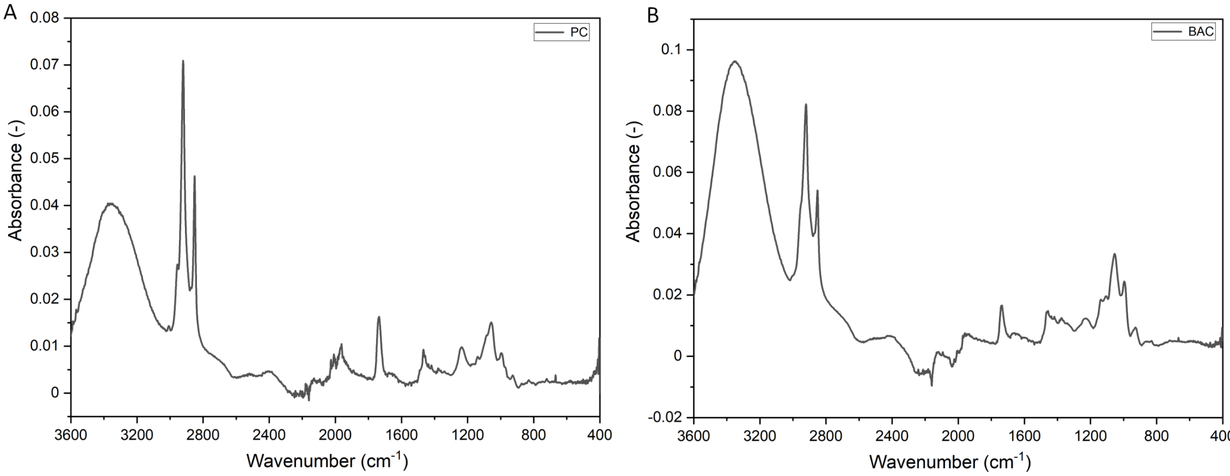


Figure S5: Raw ATR-FTIR spectra of PC (A) and BAC (B) membranes in the range of 3600–400 $cm^{-1}$.

Table S1: Tentative assignments of bands in ATR-FTIR spectra of the studied lipid model membranes in dry-film form at 22°C, where ↓ denotes a shift to a lower wavenumber compared to the POPC, and ↑ indicates a shift to a higher wavenumber compared to the PC reference.

| Group | PC | BAC | Assignment |
|---|---|---|---|
| $\nu_{as}(CH_3)$ | 2957 | 2955 ↓ | Asymmetric $CH_3$ stretching |
| $\nu_{as}(CH_2)$ | 2920 | 2917 ↓ | Asymmetric $CH_2$ stretching |
| $\nu_{as}(CH_3)$ | 2873 | 2871 ↓ | Symmetric $CH_3$ stretching |
| $\nu_{as}(CH_2)$ | 2851 | 2849 ↓ | Symmetric $CH_2$ stretching |
| $\delta(CH_2/CH_3)$ | 1468 | 1466 ↓ | $CH_2$ scissoring / $CH_3$ asymmetric deformation |
| $\delta(CH_2/CH_3)$ shoulder | 1456 | 1455 ↓ | $CH_2$ scissoring / $CH_3$ asymmetric deformation shoulder |
| $\rho(CH_2)$ | 722 | 721 ↓ | $CH_2$ rocking of hydrocarbon chains |

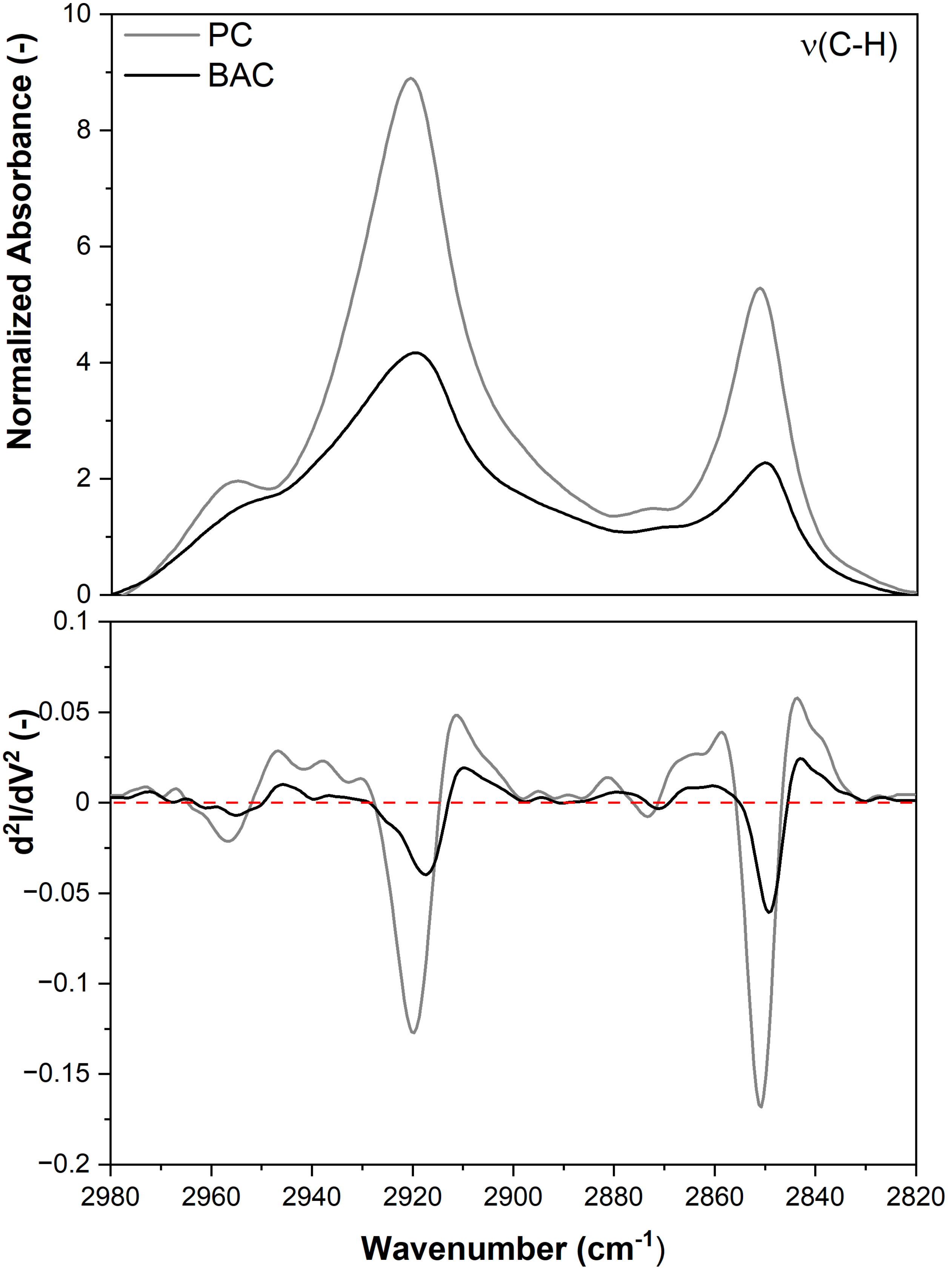


Figure S6: Normalized ATR-FTIR spectra of studied model lipid membranes PC and BAC, in the range of 2980-2820 $cm^{-1}$ with second derivatives. Spectra normalized to the $CH_2$ group (1450 $cm^{-1}$).

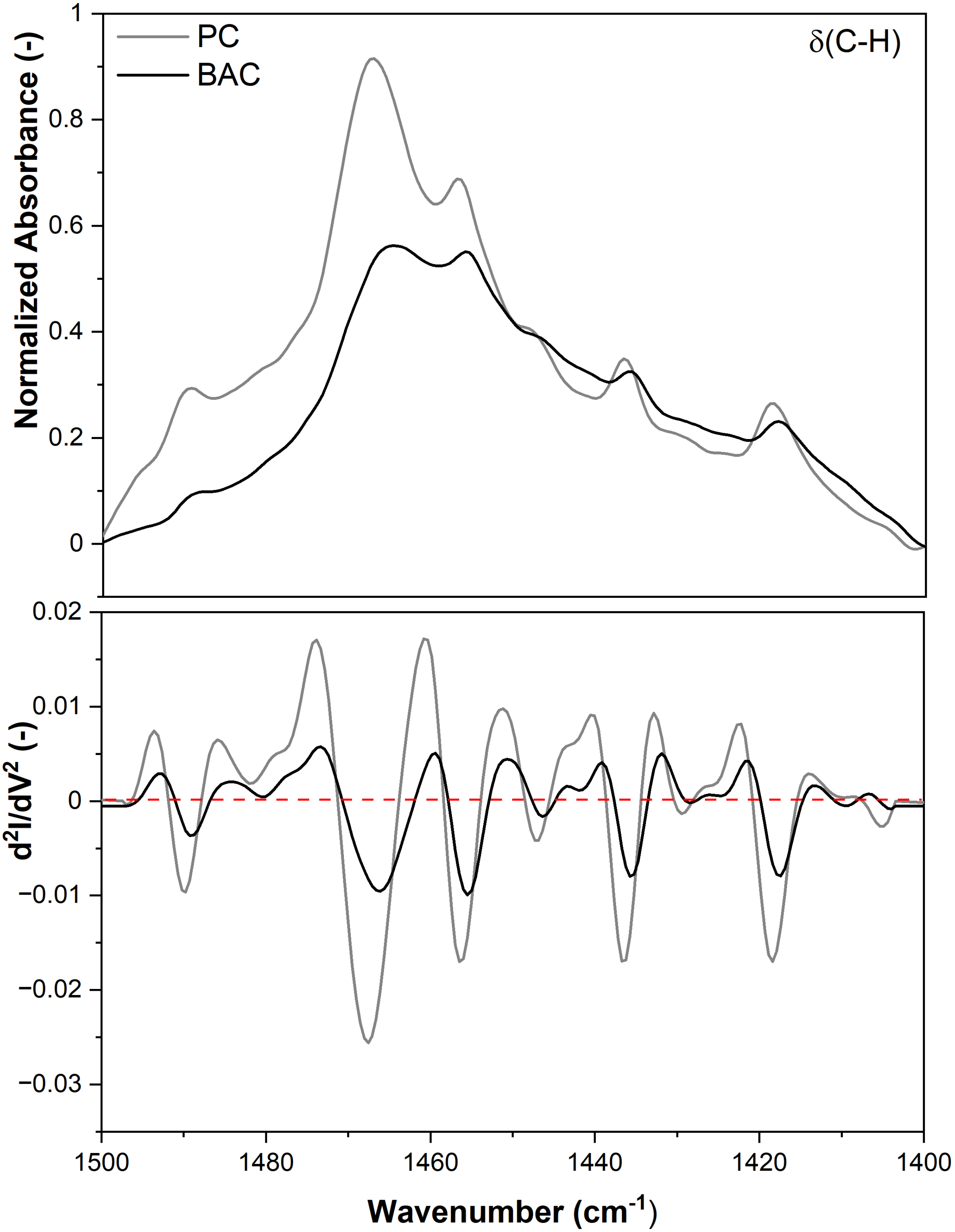


Figure S7: Normalized ATR-FTIR spectra of studied model lipid membranes POPC and BAC, in the range of 1500-1400 $cm^{-1}$ with second derivatives. Spectra normalized to the $CH_2$ group (1450 $cm^{-1}$).

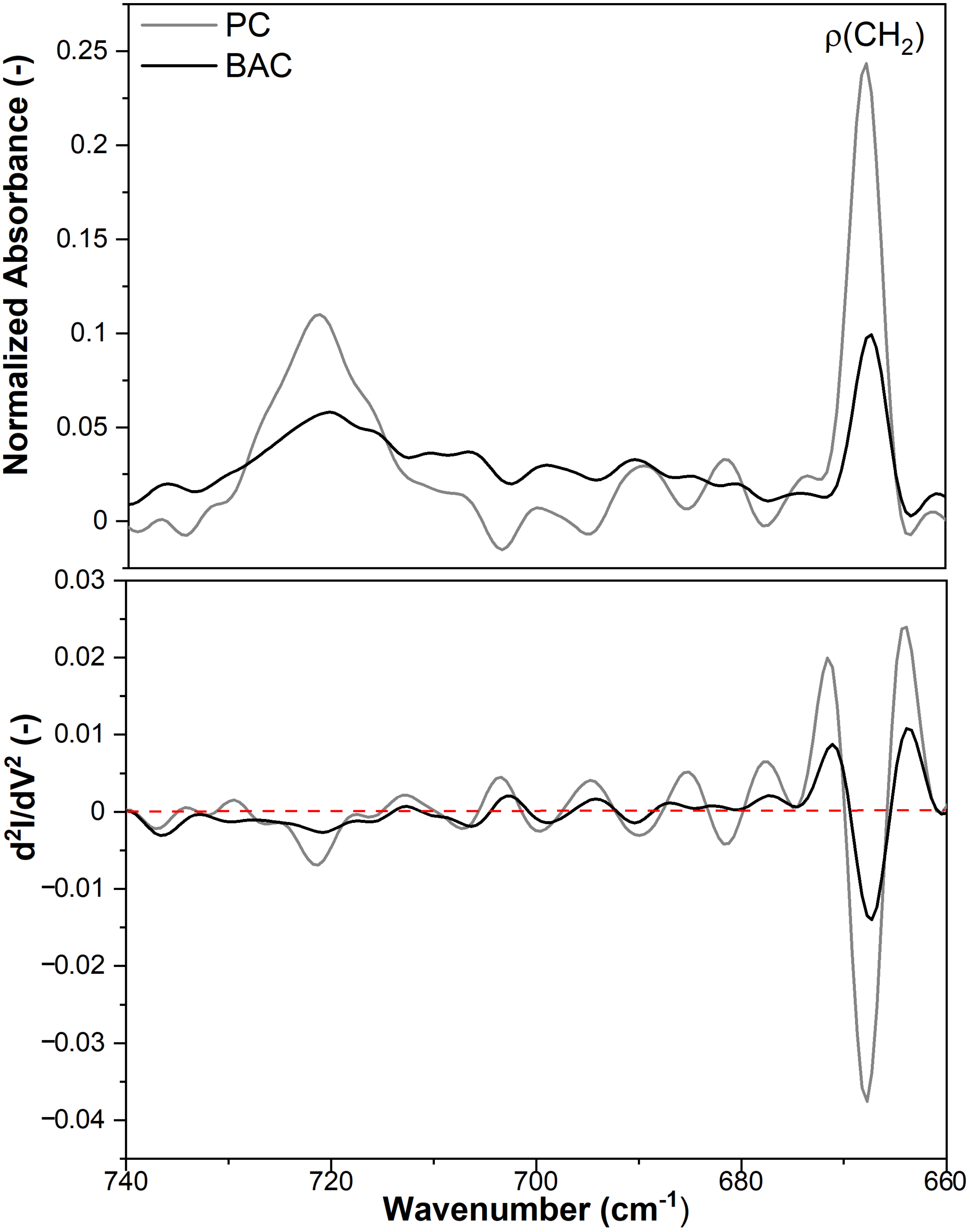


Figure S8: Normalized ATR-FTIR spectra of studied model lipid membranes POPC and BAC, in the range of 750-650 $cm^{-1}$ with second derivatives. Spectra normalized to the $CH_2$ group (1450 $cm^{-1}$).
